# Polycomb-Mediated Chromatin Loops Revealed by a Sub-Kilobase Resolution Chromatin Interaction Map

**DOI:** 10.1101/099804

**Authors:** Kyle P. Eagen, Erez Lieberman Aiden, Roger D. Kornberg

## Abstract

The locations of chromatin loops in *Drosophila* were determined by Hi-C (chemical cross-linking, restriction digestion, ligation, and high-throughput DNA sequencing). Whereas most loop boundaries or “anchors” are associated with CTCF protein in mammals, loop anchors in *Drosophila* were found most often in association with the polycomb group (PcG) protein Polycomb (Pc), a subunit of Polycomb Repressive Complex 1 (PRC1). Loops were frequently located within domains of PcG-repressed chromatin. Promoters located at PRC1 loop anchors regulate some of the most important developmental genes and are less likely to be expressed than those not at PRC1 loop anchors. Although DNA looping has most commonly been associated with enhancer-promoter communication, our results indicate that loops are also associated with gene repression.

## INTRODUCTION

Active and inactive genes are folded differently and located in different regions of the nucleus, but the molecular basis of chromosome folding and nuclear architecture remains to be determined. One folding paradigm is the formation of protein-mediated DNA loops, which are most commonly associated with enhancer-promoter communication (1). On the other hand, gene repression is most often associated with heterochromatin formation and chromosome condensation.

Chromosome folding can be revealed by chemical cross-linking, followed by restriction digestion, ligation, and high-throughput DNA sequencing (2). Such “Hi-C” analysis has revealed intra-chromosomal folding on multiple length scales. From lengths of 1 kb, the smallest so far examined (3), to hundreds of kb, two features are observed, loops (3) and so-called topologically associating domains, or “TADs” (which we and others have also referred to as A/B domains, physical domains, topological domains or contact domains) (3-7). Loops bring a pair of loci into close physical proximity; TADs represent genomic intervals in which all pairs of loci exhibit an enhanced frequency of contact, and correspond to stably condensed chromosomal regions (8). On a larger scale, extending to whole chromosomes, TADs interact with one another to form “compartments” (2).

Hi-C on the smallest length scale, based on the most extensive sequencing, is needed for unambiguous identification of chromatin loops, and has been reported thus far only for the mouse and human genomes (3). We have now overcome this limitation in *Drosophila*, by Hi-C analysis at sub-kilobase resolution. We find an unanticipated correlation with results of ChIP-seq analysis in *Drosophila*, with an important functional correlate.

## RESULTS

### Sub-kilobase resolution *Drosophila* Hi-C

We performed Hi-C on embryonic Kc167 *Drosophila melanogaster* cultured cells and identified 529 million chromosomal contacts (read pairs that remain after exclusion of duplicates, unligated fragments, and reads that align poorly with genome sequences). The resolution of the resulting contact maps was 260 bp, or “sub-kilobase”, based on standard metrics (3) (Fig. S1, Table S1, and Methods). For this reason, and because 77.1% of the restriction fragments used for the Hi-C analysis were shorter than 500 bp, we plotted the contact maps in units of at least 500 bp. The contact maps were 3- to 4 fold more dense than those previously reported for human cells at kilobase resolution (3).

Consistent with previously reported *Drosophila* Hi-C contact maps (4, 8, 9), we observed both TADs (apparent in a Hi-C contact map as boxes of enriched contact frequency tiling the diagonal; Fig. 1A and 1B) and compartments (apparent as off-diagonal boxes of alternating enriched or depleted contacts, whose boundaries are aligned with those of the on-diagonal boxes; Fig. 1a,b and Table S2). We identified more TADs in Kc cells than were originally reported (9) (2,126 versus 1,110; Table S2) because more TAD boundaries could be detected at higher resolution (3). We also identified chromatin loops, which appeared as focal peaks of contact enrichment (Fig. 1C and Table S3), and which were previously unobservable in lower resolution *Drosophila* maps.

**Fig. 1.**
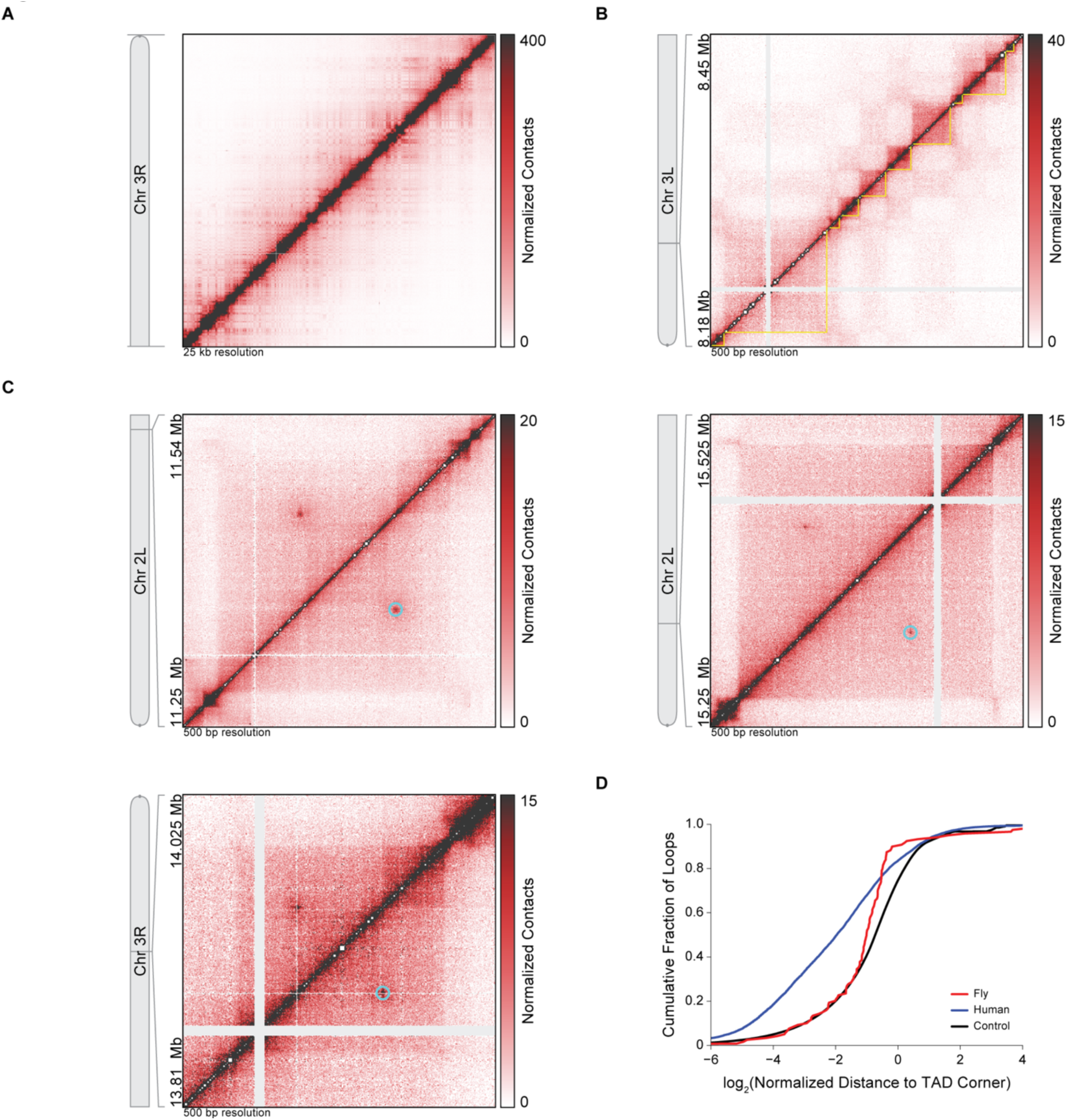
Sub-kilobase resolution Hi-C identifies *Drosophila* chromatin loops. **(A)** Intra-chromosome 3R Hi-C contact map at 25 kb resolution shows off-diagonal boxes of alternating enriched or depleted contacts indicative of compartments. **(B)** Hi-C contact map at 500 bp resolution of a region of chromosome 3L shows TADs (yellow outlines) as boxes of enriched contact frequency tiling the diagonal. On-diagonal boxes (TADs) align with off-diagonal boxes indicative of compartmentation. **(C)** Hi-C contact maps at 500 bp resolution reveal the presence of chromatin loops, identified as focal peaks of contact enrichment (cyan circles). **(D)** Cumulative distributions of the two-dimensional Euclidean distance between a loop and the closest TAD corner for *Drosophila* loops (red), human loops (blue), and the mean of 10,000 shuffled sets of *Drosophila* loops (black) with 95% confidence interval (gray shading). *Drosophila* loops are farther from TAD corners than human loops (*P* = 3.79 x 10^−11^; two-sided KS test), but closer to TAD corners than the shuffled control (*P* = 5.89 x 10^−9^; two-sided KS test). Distance is normalized by TAD size.

### *Drosophila* Loops Are Unrelated to TADs

We identified 120 chromatin loops, far fewer than the number of TADs, and also far fewer than the 9,448 loops identified in human GM12878 B-lymphoblastoid cells (3), even after accounting for the difference in genome size.

*Drosophila* loops differed from mammalian loops in their relationship to TADs (Fig. 1D). Mammalian loop anchors are frequently (38% of loops) located at TAD corners, and therefore coincide with TAD boundaries. In contrast, the vast majority (82.5%) of *Drosophila* loops did not appear at TAD corners (Fig. 1C), and conversely, 99.1% of *Drosophila* TADs did not have focal peaks at their corners. Evidently, chromatin condensation revealed by TADs (8) is not due in *Drosophila* to loops detectable by our methods.

### Lack of CTCF at *Drosophila* Loop Anchors

In humans, 86% of loop anchors are associated with CTCF, and the CTCF-binding motifs are in a convergent orientation (3, 10). In contrast, *Drosophila* loop anchors did not tend to align with CTCF ChIP-seq signals (11) (Fig. 2A and 2B), and only 28.2% of loop anchors overlapped CTCF-binding sites genome-wide (Fig. 2C). Loop anchors were less likely to occur at the strongest CTCF ChIP-seq peaks than at the weakest peaks (Fig. 2C, left). Conversely, the strongest CTCF ChIP-seq peaks were less likely to occur at loop anchors than at the weakest peaks (Fig. 2C, right). At a minimum ChIP enrichment of 16-fold, loop anchors are 9.55-fold enriched at CTCF sites, but this accounts for only 13.2% of loop anchors and 1.66% of CTCF sites. Evidently, CTCF is rarely, if ever, significantly associated with chromatin loops, and the vast majority of CTCF sites are unrelated to the chromatin loops we identify in *Drosophila.* Similar results were obtained for the other *Drosophila* insulator proteins BEAF-32, Su(Hw), and CP190 (Fig. S2). In contrast to the results for CTCF, and in keeping with reports for human loop anchors, the majority (72.8%) of *Drosophila* loop anchors were associated with the cohesin subunit Rad21. Moreover, stronger Rad21 ChIP peaks were more likely to overlap with loop anchors (Fig. 2A, 2B, and 2D).

**Fig. 2.**
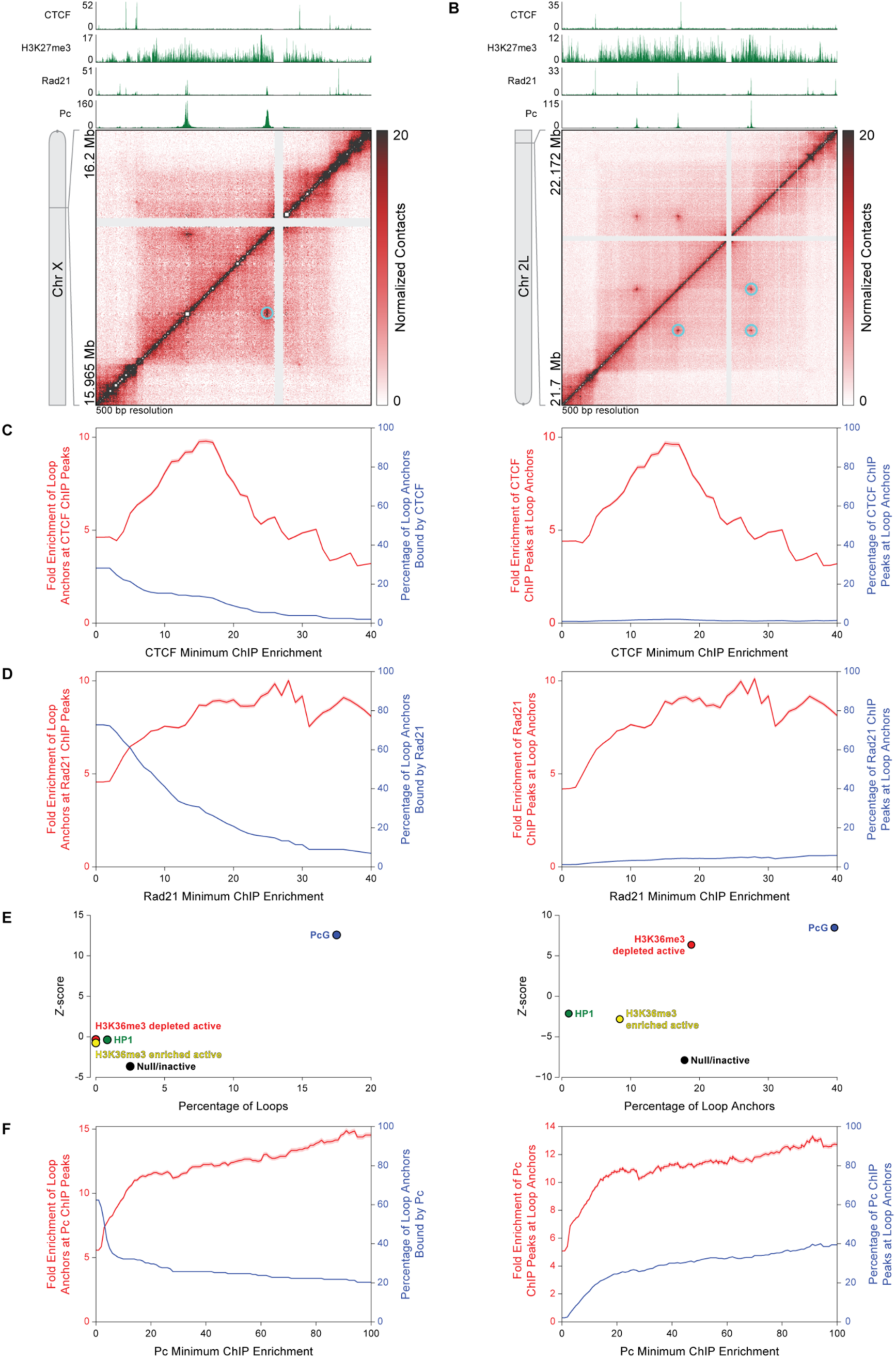
*Drosophila* loop anchors are bound by PRC1. **(A)** Hi-C contact map at 500 bp resolution of a region of chromosome X shows chromatin loops (cyan circles) that align with *Pc* ChIP-seq peaks. CTCF, histone H3K27me3, Rad21, and *Pc* ChIP-seq profiles are aligned above the map. **(B)** Hi-C contact map at 500 bp resolution of a region of chromosome 2L shows a small network of chromatin loops (cyan circles) that align with *Pc* ChIP-seq peaks. CTCF, histone H3K27me3, Rad21, and *Pc* ChIP-seq profiles are aligned above the map. **(C)** Left, fold enrichment of loop anchors at CTCF ChIP peaks (red) and percentage of loop anchors bound by CTCF (blue) at the CTCF minimum ChIP enrichment indicated on the abscissa. Right, fold enrichment of CTCF ChIP peaks at loop anchors (red) and percentage of CTCF ChIP peaks at loop anchors (blue) at the CTCF minimum ChIP enrichment indicated on the abscissa. Red shading indicates 95% confidence interval. **(D)** Same as **(C)** except for Rad21. **(E)** Left, Z-score of the enrichment/depletion of entire loops within five functional classes (“colors”) of chromatin relative to 10,000 random shuffle controls shows that entire loops are strongly enriched in PcG-repressed chromatin. Right, Z-score of enrichment/depletion of entire loop anchors within five functional classes (“colors”) of chromatin relative to 10,000 random shuffle controls shows that entire loop anchors are strongly enriched in PcG-repressed chromatin. Enrichment is given by a positive Z-score, depletion by a negative Z-score. **(F)** Same as **(C)** except for *Pc.*

### Loops are Found within Polycomb-Repressed Chromatin

For a more comprehensive assessment of the relationship between loops, loop anchors, and chromatin-bound proteins, we compared our loop annotation to the five functional classes of chromatin that were previously identified on the basis of genome-wide non-histone protein localization in *Drosophila* Kc cells: PcG-repressed chromatin, HP1 heterochromatin, another type of repressed chromatin (“null/inactive”), and two types of active chromatin (“H3K36me3-enriched or –depleted”) (12). We found that 17.5% of loops were entirely contained within a region of PcG-repressed chromatin, and such enrichment of entire loops within PcG-repressed chromatin was far greater than expected on a random basis (10.9-fold, *P* = 1.82 x 10^−35^) (Fig. 2E). Few loops were entirely located within H3K36me3-enriched or -depleted active chromatin (no loops), or HP1 heterochromatin (0.8% of loops), and loops were neither significantly enriched nor depleted from these classes (Fig. 2E). Loops were depleted from null/inactive chromatin (2.5% of loops, a 5.1-fold depletion, *P* = 5.07 x 10^−4^). The low percentage of loops entirely located within a single functional chromatin class indicates that the majority of loops must span more than one class.

Many loop anchors (39.6%) were located within PcG-repressed chromatin, and enrichment of loop anchors within PcG-repressed chromatin was greater than expected on a random basis (2.35-fold, *P* = 1.79 x 10^−16^) (Fig. 2E). A smaller fraction of loop anchors was found in H3K36me3-depleted active chromatin (18.8% of anchors, 2.76-fold enriched, *P* = 6.01 x 10^−10^) (Fig. 2E). Loop anchors were significantly depleted from HP1 heterochromatin, null/inactive chromatin, and H3K36me3-enriched active chromatin (HP1: 0.1% of anchors, 3.32-fold depletion, *P* = 0.0436; null/inactive: 17.8%, 2.47-fold depletion, *P* = 1.19 x 10^−14^; H3K36me3-enriched active: 8.42%, 1.82-fold depletion, *P* = 7.07 x 10^−3^) (Fig. 2E).

### The Polycomb Protein is Found at Loop Anchors

PcG proteins are involved in gene repression during development and are components of two evolutionarily conserved complexes, PRC1 and PRC2. PRC2 trimethylates histone H3 at lysine 27, which demarcates PcG-repressed chromatin and is bound by the chromodomain of the Polycomb (Pc) subunit of PRC1 (13). Loops were readily identified within H3K27me3-enriched regions (14) (Fig. 2A, 2B, 3A, and 3B), and loop anchors tended to coincide with Pc ChIP-seq signals (15) from Kc cells (Fig. 2A, 2B, 3A, and 3B). Occasionally, loop anchors that aligned with Pc ChIP signals formed networks of interactions (Fig. 2B). Overall, a remarkable 62.4% of loop anchors were associated with Pc ChIP peaks. The enrichment of loop anchors at Pc ChIP peaks increased with the strength of the Pc ChIP peak, and enrichment of Pc ChIP peaks at loop anchors continually increased with the strength of the Pc ChIP peak as well (Fig. 2F). The percentage of Pc ChIP peaks at loop anchors increased 18.8-fold from 2.1% to 39.5% as the strength of the Pc peak increased from 0-fold to 100-fold minimum ChIP enrichment. This is in contrast with CTCF, BEAF-32, Su(Hw), and CP190, none of which exhibited a monotonic relationship between peak strength and the frequency of overlap with loop anchors (Fig. 2C and S2). To ensure that these results reflected the presence of Pc, rather than off-target binding by the antibody that was employed for ChIP, we analyzed ChIP-seq data from a second antibody against Pc (15) and obtained essentially the same results (Fig. S3).

**Fig. 3.**
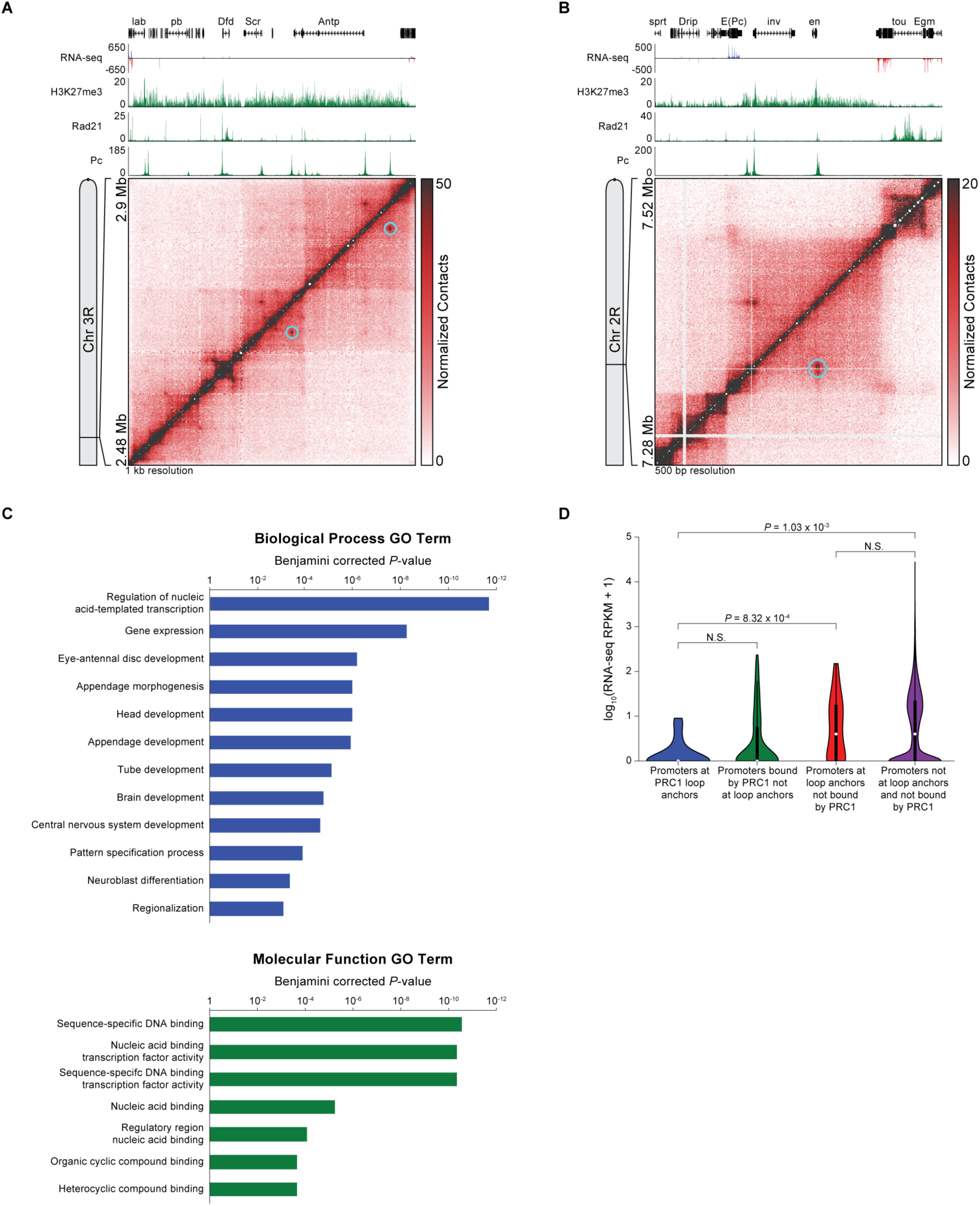
Developmentally regulated promoters are found at PRC1 loop anchors. **(A)** Hi-C contact map at 1 kb resolution shows chromatin loops (cyan circles) at the ANT-C Hox gene complex. RNA-seq, histone H3K27me3, Rad21, and *Pc* ChIP-Seq profiles are aligned above the map. **(B)** Hi-C contact map at 500 bp resolution shows chromatin loops (cyan circles) at the *inv* and *en* promoters. RNA-seq, histone H3K27me3, Rad21, and *Pc* ChIP-Seq profiles are aligned above the map. **(C)** GO term P-value chart indicates that PRC1 loop-anchor promoters are enriched for genes that regulate transcription and development. **(D)** Promoters a PRC1 loop anchors are less likely to be expressed than promoters at loop anchors not bound by PRC1 (*P* = 8.34 x 10^−4^; one-sided Mann-Whitney U-test) or promoters not at loop anchors and not bound by PRC1 (*P* = 1.03 x 10^−3^; one-sided Mann–Whitney *U*-test). Expression levels indicated by violin plots that are overlaid with Tukey box plots (black) and the median expression level (white circle). N.S., not significant.

The presence of strong Pc ChIP sites that are not associated with loop anchors may reflect the presence of chromatin loops that we cannot detect. Especially in the case of strong Pc ChIP sites separated by only a few kilobases, it is difficult to identify focal peaks in the Hi-C map due to the strong signal along the diagonal.

### PRC1 Loops Are Associated with Gene Repression

We found that Pc ChIP sites enriched at least 30-fold were not only particularly large (in terms of 1D extent along the DNA sequence, see for instance Pc ChIP track in Fig. 2A and 2B), but the majority of these sites (64.2% versus 21.8% for random shuffle control) contained both GAGA and PHO motifs, which are characteristic of polycomb response elements bound by PRC1 (16). We therefore refer to the anchors found at such sites as “PRC1 loop anchors”. Loop anchors were 11.5-fold enriched relative to random shuffle control at these sites, which accounted for 26% of all loop anchors and 26.2% of Pc sites.

A large fraction (32.7%) of PRC1 loop anchors were found at promoters of important developmentally regulated genes, including those for the homeotic PcG target genes *Antennapedia (Antp)* and *Sex combs reduced (Scr)* at the Antennapedia Hox gene complex (ANT-C) (Fig. 3A), as well as *invected (inv)* and *engrailed (en)* (Fig. 3B). Gene ontology (GO) term analysis indicated that genes involved in regulating transcription and development were particularly enriched at PRC1 loop-anchor promoters (Fig. 3C and Table S4).

PRC1 loop-anchor promoters were less likely to be expressed than promoters at loop anchors not bound by PRC1 (0 compared to 3 median RPKM expression level, *P* = 8.32 x 10^−4^; one-sided Mann–Whitney *U*-test) or promoters not bound by PRC1 and not at loop anchors (0 compared to 3 median RPKM expression level, *P* = 1.03 x 10^−3^; one-sided Mann–Whitney *U*-test) (Fig. 3D). Maximum expression levels of promoters at PRC1 loop anchors were less than that of promoters bound by PRC1 but not at loop anchors, even though median expression levels did not significantly differ. A similar result was obtained for promoters at loop anchors not bound by PRC1 compared to promoters not at loop anchors and not bound by PRC1 (Fig. 3D). Taken together, these results suggest a strong relationship between gene repression and chromatin looping, but whether loops cause gene repression or form as a consequence of gene inactivity remains to be determined.

## DISCUSSION

The locations of loop anchors in *Drosophila* determined here are notable both for correlations with ChIP-seq data and for the lack thereof. The lack of correlation with locations of CTCF protein was unexpected, inasmuch as most loop anchors in mammals are associated with CTCF protein, apparently bound to CTCF sequence motifs in a convergent orientation (3, 10). There are evidently multiple patterns of protein association with loop anchors in metazoans. The association of loop anchors in *Drosophila* with Pc protein is noteworthy because it points to a role of looping not only in gene activation, as widely observed in the past, but in gene repression as well.

Regions of PcG-repressed chromatin (“PcG domains”) that are separated by hundreds of kilobases to megabases are known to be in enhanced spatial proximity (4, 17, 18), but details of their internal organization have only been investigated by averaging over many PcG domains (16). The high resolution of our Hi-C contact maps revealed chromatin loops within individual PcG domains, giving insight into their internal organization. PRC1 is known to compact nucleosome arrays *in vitro* (19). Knockdown of the PRC1 subunit Polyhomeotic (Ph) *in vivo* decompacts PcG-repressed chromatin(20), and Ph that is unable to polymerize impairs the ability of PRC1 to form cluster (21). Together with these findings, our results suggest that PRC1-bound chromatin loops within PcG-repressed domains either establish or maintain a condensed state.

Previous analyses by 3C have pointed to associations of PcG proteins with chromatin loops for the Bithorax complex (BX-C) in S2 cells (22), for *inv* and *en* in BG3 and Sg4 cells (23), and for an embryonic, pupae, and adult transgenic reporter system (24). Our Hi-C data are, however, at higher resolution and genome-wide. Higher resolution allowed more comprehensive analysis, such as the unambiguous identification of loops and the segmentation of ANT-C into a series of TADs with one or two homeotic gene promoters per TAD (Fig. S4). Genome-wide analysis revealed both the pervasive nature of Pc protein association and the absence of significant CTCF protein association, despite conservation of CTCF from *Drosophila* to human (25).

Another report on *Drosophila* chromatin loops in Kc167 cells appeared while our manuscript was in preparation (26). The other authors noted an enrichment of cohesin but a lack of *Drosophila* CTCF at loop anchors, consistent with our observations. The other authors did not mention Polycomb, but our analysis of their data revealed an enrichment of loop anchors at Pc ChIP peaks and an enrichment of Pc ChIP peaks at loop anchors (Fig. S5). We find a likelihood of repression of promoters at Pc-bound loop anchors, especially for developmental genes; the other authors observed an enrichment of active developmental enhancers at loop anchors, possibly because these are among the many loop anchors not bound by Pc, or because the chromatin at these loop anchors is bivalent, bound by non-histone proteins and histone post-translational modifications associated with both gene activation and repression (27).

The occurrence of PRC1 at loop anchors could reflect a role in loop formation similar to that proposed for CTCF in mammals, wherein cohesin-complexes extrude loops, in a process halted upon reaching bound CTCF (10, 28, 29). Consistent with this model, a large majority (72.8%) of Drosophila loop anchors are bound by the Rad21 subunit of cohesin. Regardless of whether PRC1 performs such a role, additional proteins must be involved, because PRC1 is present at only 26% of *Drosophila* loop anchors.

## MATERIALS AND METHODS

### Hi-C

Hi-C was performed using the tethered conformation capture approach (8, 30), which improves the signal-to-noise ratio needed for detection of chromatin loops. In brief, *Drosophila melanogaster* Kc167 cultured cells were fixed with 1% EM grade paraformaldehyde. Cells were lysed and cross-linked proteins were biotinylated at cysteine residues and the DNA digested with DpnII. Digested chromatin was bound to streptavidin beads, thoroughly washed to remove uncross-linked DNA, DNA ends filled in with biotin-14-dATP, and free DNA ends ligated together. DNA-protein cross-links were reversed, DNA purified, biotinylated nucleotides marking unligated ends removed, and the DNA sheared to approximately 500 bp. The biotinylated DNA was pulled down with streptavidin beads, prepared for and subjected to high-throughput Illumina DNA sequencing. Further details provided in *SI Appendix, Supplementary Materials and Methods*.

### Hi-C Analysis

Hi-C data was analyzed using the Juicer pipeline as previously described (3, 31). In brief, Hi-C reads were mapped to the dm3 reference genome using BWA-MEM. Aligned reads were assigned to restriction fragments, duplicates removed, reads with a MAPQ < 30 removed, and intra-fragment reads removed. The genome was then divided into equally spaced bins and the number of contacts was counted in each pair of bins. Hi-C contact maps were normalized by matrix balancing (3, 31).

TADs were identified using the previously described Arrowhead algorithm (3, 31) and loops were identified by visual inspection and manual annotation using Juicebox (32). Further details provided in *SI Appendix, Supplementary Materials and Methods*.

External datasets used in this study can be found in Table S5.

## AUTHOR CONTRIBUTIONS

K.P.E. conceived the project, designed the study, performed Hi-C experiments, and analyzed the data. All authors discussed and interpreted the results. K.P.E. and R.D.K. wrote the manuscript with comments from E.L.A.

The authors declare no conflict of interest.

Data deposition: The Hi-C sequence data reported in this paper have been deposited at the GEO database, www.ncbi.nlm.nih.gov/geo (accession no. GSE89112).

## ACKNOWLEDGEMENTS

We are grateful to Suhas S.P. Rao for suggestions regarding loop annotation and Su-Chen Huang and Olga Dudchenko for assistance with DNA sequencing. This research was supported by NIH grants GM36659 and AI21144 to R.D.K. and by NIH New Innovator Award 1DP2OD008540, NIH 4D Nucleome Grant U01HL130010, NSF Physics Frontier Center PHY-1427654, Welch Foundation Q-1866, Cancer Prevention Research Institute of Texas Scholar Award R1304, an NVIDIA Research Center Award, an IBM University Challenge Award, a Google Research Award, a McNair Medical Institute Scholar Award, and the President’s Early Career Award in Science and Engineering to E.L.A.

## APPENDIX A SI APPENDIX

## SUPPLEMENTARY MATERIALS AND METHODS

### Cell Culture

Kc167 cells were obtained from the Drosophila Genomics Resource Center (Bloomington, IN). Cells were grown in CCM3 media (HyClone) at 25° C.

### Hi-C Library Preparation

Hi-C libraries were prepared using a modification of the previously described tethered conformation capture (TCC) (8, 30) protocol. 1 billion Kc167 cells at a density of 4-6 x 10^6^ cells/mL were washed with 20 mL CMM3 media and then cross-linked with 1% EM-grade paraformaldehyde in 450 mL CCM3 media for 10 minutes at room temperature while mixing. Paraformaldehyde was quenched by adding 29.8 mL 2.5 M glycine (150 mM final) and incubating for 5 minutes at room temperature while mixing. Cells were washed with 10 mL ice-cold PBS, pelleted, flash frozen in liquid nitrogen and stored at -80°C.

Cells were thawed, resuspended in 10 mL lysis buffer (10 mM Tris-HCl pH 8.0, 10 mM NaCl, 0.2% Igepal CA-630, 1 mM PMSF, 2 mM bezamidine, 2 μM pepstatin A, 0.6 μM leupeptin), and then incubated on ice for 15 minutes. Cells were transferred to a 15 mL Dounce homogenizer and lysed by applying 15 strokes of pestle B, cooled on ice for 1 minute, followed by applying another 15 strokes of pestle B. The lysate was centrifuged at 2,500 x g for 5 minutes, the pellet washed with 5 mL lysis buffer, centrifuged again at 2,500 x g for 5 minutes, the pellet resuspended in 4.6 mL of wash buffer 1 (50 mM Bis-Tris-HCl pH 6.0, 100 mM NaCl, 10 mM MgCl2, 0.1% SDS), incubated at 65° C for 10 minutes, and then immediately cooled on ice.

1.4 mL 25 mM EZ-link Iodoacetyl-PEG2-Biotin (IPB; in wash buffer 1; Thermo Scientific) was added to the sample and incubated for 1 hour at room temperature while rotating. 600 μL of 10% Triton X-100 was added, gently mixed, then 160 μL of 50 U/μL DpnII (NEB) was added and the sample incubated at 37° C for 2 hours while rotating. 1.298 mL 10% SDS was added, incubated at 65° C for 30 minutes, and then immediately cooled on ice. The sample was added to a 20 kD MWCO, 3-12 mL Slide-A-Lyzer Dialysis Cassette (Thermo Scientific) and dialyzed at room temperature against 4 L of dialysis buffer (10 mM Tris-HCl pH 8.0, 1 mM EDTA). After 2 hours the buffer was replaced with 4 L of fresh dialysis buffer and dialysis continued overnight.

800 μL of Dynabeads MyOne Streptavidin T1 beads (Life Technologies) were washed three times with 2 mL PBST (PBS + 0.01% Tween 20) and then resuspended in 1 mL PBST. The dialyzed sample was rapidly added to the beads and incubated at room temperature for 30 minutes while rotating. 50 μL of 25 mM neutralized IPB (treated with 10-fold excess β-mercaptoethanol) was added to the sample and then incubated at room temperature for 15 minutes while rotating.

Beads were washed once with 3 mL PBST, twice with 3 mL wash buffer 2 (10 mM Tris-HCl pH 8.0, 50 mM NaCl, 0.4% Triton X-100), resuspended in 1 mL of wash buffer 2, and divided across 5 equal volume aliquots. 190 μL of a fill-in master mix (654 μL water, 11 μL 1 M MgCl_2_, 110 μL NEBuffer 2, 7.7 μL 10 mM dTTP, 7.7 μL 10 mM dCTP, 7.7 μL 10 mM dGTPαS, 192.5 μL 0.4 mM Biotin-14-dATP, 44 μL 10% Triton X-100) was added to each aliquot followed by 10 μL of 5 U/μL DNA Polymerase I, Large (Klenow) Fragment (NEB) and then incubated at 37° C for 75 minutes while rotating. The reaction was stopped with 10 μL 0.5 M EDTA.

Each aliquot of beads was washed twice with 500 μL wash buffer 3 (50 mM Tris-HCl pH 7.4, 0.4% Triton X-100, 0.1 mM EDTA) and then resuspended in 500 μL of wash buffer 3. 8.99 mL of a ligation master mix (37.972 mL water, 5.179 mL 10% Triton X-100, 5.179 mL 500 mM Tris-HCl pH 7.5, 100 mM MgCl_2_, 556 μL 10 mg/mL BSA, 556 μL 100 mM ATP, 556 μL 1 M DTT) was added to each aliquot followed by 10 μL 2,000 U/μL T4 DNA Ligase (NEB) and then incubated at room temperature for 4 hours while rotating. The reaction was stopped with 400 μL 0.5 M EDTA.

Beads were collected with a magnet, the supernatant discarded, and the beads resuspended in 300 μL of extraction buffer (50 mM Tris-HCl pH 8.0, 100 mM NaCl, 1 mM EDTA, 0.2% SDS). 25 μL 800 U/mL proteinase K (NEB) was added to each aliquot and each aliquot was incubated at 65° C overnight.

Beads were collected with a magnet and the supernatant transferred to a fresh tube. The DNA was precipitated with ethanol, washed twice with 70% ethanol, and all aliquots combined and resuspended in a total volume of 75 μL 10 mM Tris-HCl pH 8.0. The sample was incubated at 42° C for 15 minutes followed by addition of 5 μL 1 mg/mL RNase A and incubation at 37° C for 30 minutes. DNA concentration was determined with a Qubit dsDNA HS Assay.

7 μL water, 10 μL 10x NEBuffer 1, 1 μL 10 mg/mL BSA, and 3 μL 100 U/μL exonuclease III was added to 79 μL of DNA (no more than 10 μg DNA per reaction, if more than 10 μg multiple reactions were performed in parallel), and incubated at 37° C for 1 hour. The reaction was stopped by adding 2 μL 0.5 M EDTA, 2 μL 5 M NaCl, and incubating at 70° C for 20 minutes. Total sample volume was adjusted to 130 μL.

DNA was sheared to 500 bp with a Covaris S2 at duty cycle 10%, intensity 4, 200 cycles/burst for 55 seconds. 125 μL of sample was transferred to a fresh tube and the DNA was size-selected by first adding 68.8 μL (0.55x volumes) of SPRIselect beads (Beckman), vortexed to mix, and incubated at room temperature for 5 minutes. Beads were collected with a magnet, the supernatant transfered to a fresh tube. 25 μL of SPRIselect beads were added to the supernatant, vortexed to mix, and incubated at room temperature for 5 minutes. Beads were collected with a magnet, the supernatant discarded, and, while still on the magnet, the beads were washed twice with 200 μL 85% ethanol and then air dried. DNA was eluted by resuspending the beads in 53 μL of 10 mM Tris-HCl pH 8.0 and incubated at room temperature for 5 minutes. Beads were collected with a magnet, and 51 μL of the eluate was transferred to a fresh tube. DNA concentration was determined with a Qubit dsDNA HS Assay.

DNA was initially prepared for high-throughput sequencing following the directions for “NEBNext End Prep” and “Adaptor Ligation” in the NEBNext Ultra II DNA Library Prep Kit for Illumina (NEB) (if more than 1 μg of DNA, multiple reactions were performed in parallel). 50 μL of Dynabeads MyOne Streptavidin C1 beads (Life Technologies) were washed twice with 100 μL 1x B&W buffer + 0.05% Tween 20 (2x Bind & Wash (B&W) buffer: 10 mM Tris-HCl pH 7.5, 2 M NaCl, 1 mM EDTA) and then resuspended in 86.5 μL of 2x B&W buffer. Adaptor-ligated DNA was added to the beads and incubated at room temperature for 30 minutes while rotating. Beads were collected with a magnet, the supernatant discarded, the beads washed twice with 100 μL 1x B&W buffer + 0.05% Tween 20, and then twice with 100 μL 10 mM Tris-HCl pH 8.0, 1 mM EDTA.

DNA was eluted by resuspending the beads in 5 μL of 95% freshly deionized formamide, 10 mM EDTA pH 8.0, and incubating at 90° C for 10 minutes. The beads were then cooled to room temperature and collected with a magnet. The supernatant was transferred to a fresh tube and mixed with 15 μL 10 mM Tris-HCl pH 8.0.

The optimal number of PCR cycles for library amplification was determined by setting up an analytical qPCR reaction: 1 μL water, 1 μL eluted DNA, 1 μL primer mix (2.5 μM NEB Universal PCR Primer, 2.5 μM NEB Index Primer, 2.5 mM MgCl_2_, 5x SYBR Green I, 5x ROX), 2.5 μL NEBNext Ultra II Q5 Master Mix. The sample was thermocycled on a qPCR machine at 98° C for 30 seconds followed by 25 cycles of 98° C for 10 seconds, 65° C for 75 seconds. The linear Rn versus cycle number was plotted to determine the cycle number corresponding to one-third of maximum fluorescence. Final library amplification was performed by setting up the following PCR in duplicate: 10.5 μL water, 2.5 μL 10 mM MgCl_2_, 1 μL 25 μM NEB Universal PCR Primer, 1 μL 25 μM NEB Index Primer, 10 μL eluted DNA, 25 μL NEBNext Ultra II Q5 Master Mix. The sample was thermocycled at 98° C for 30 seconds followed by 3 or 4 cycles (as determined above) of 98° C for 10 seconds, 65° C for 75 seconds, followed by 65° C for 5 minutes, and then held at 4° C. The amplified library was purified as in “Cleanup of PCR Amplification” in the NEBNext Ultra II DNA Library Prep Kit for Illumina (NEB), adjusting the volumes as necessary. The DNA was eluted in 17 μL 10 mM Tris-HCl pH 8.0 and 15 μL was collected as the final eluate.

DNA concentration was determined with a Qubit dsDNA HS Assay, DNA integrity determined by a Bioanalyzer High Sensitivity DNA Chip, and then accurately quantified using a KAPA Library Quantification Kit. DNA was paired-end sequenced on an Illumina NextSeq 500 or HiSeq X instrument.

### Hi-C Data Processing

Data was processed as previously described (3, 31). In brief, reads were mapped to dm3/BDGP Release 5 of the *Drosophila melanogaster* genome using BWA-MEM, reads were assigned to restriction fragments, duplicates removed, reads with a MAPQ < 30 removed, and intra-fragment reads removed. The genome was then divided into equally spaced bins and the number of contacts was counted in each pair of bins.

We noticed at kilobase and sub-kilobase resolution many empty row/columns of the matrix due to restriction fragments spanning multiple bins. Therefore, for high resolution maps, to account for the uncertainty in the location of the position of the cross-link within each restriction fragment, we randomized the read position within the restriction fragment each read mapped to and then assigned the resulting contact to its respective genomic bin. This improved the quality of the maps as “missing data” was now “recovered” without changing the position of loops or TADs. All subsequent analysis was performed using randomized intrafragment read positions. Based on the previously established metric for Hi-C map resolution (3), the resolution of our Hi-C contact maps is 260 bp. However, this approaches the median restriction fragment length for DpnII (193 bp) and, consequently, may overestimate the true resolution. For the dm3 reference genome 77.1% of restriction fragments are shorter than 500 bp, which indicates an appropriate lower bound for map resolution while at the same time recovering high-resolution information from deeply sequenced *Drosophila* Hi-C libraries.Therefore, using the previously defined metric (3), we have attained the maximum possible resolution using DpnII for fragmentation or “sub-kilobase” resolution.

Hi-C contact maps were normalized by matrix balancing using the KR normalization algorithm (3, 31, 33). All subsequent analysis was performed on KR normalized contact maps. Reproducibility between biological replicates was determined by flattening the Hi-C matrices to vectors and calculating the Pearson's correlation coefficient (*r*) between the vectors. Biological replicates were highly correlated at all resolutions (Extended Data Fig. 1D). Therefore, datasets were merged after filtering, intrafragment read positions randomized as described above, and the combined contact map KR normalized. All subsequent analysis was performed on the combined, KR normalized contact map.

TADs were identified using the previously described Arrowhead algorithm (3, 31), except with the addition of a post-processing step. This step was necessary as the Arrowhead algorithm applied at 500 bp resolution skipped some obvious larger domains, whereas the Arrowhead algorithm applied at 5 kb resolution cannot identify smaller domains. Therefore, TADs were identified at 5 kb, 2 kb, 1 kb, and 500 bp resolution using the Arrowhead algorithm, merged into a single list sorted by decreasing corner score, and conflicts, defined as the boundary of one TAD being located within another TAD, resolved by using the greater corner score of any conflicting TADs. Use of the greater corner score ensures that the most prominent, high confidence TADs are identified. Although this procedure precludes the identification of nested TADs, visual inspection of the resulting TAD annotation revealed excellent agreement with Hi-C contact maps and is consistent with prior annotations of non-nested TADs in *Drosophila* (4, 9).

*Drosophila* loops were manually annotated by visual identification of focal peaks of contact enrichment using Juicebox (32). The number of Hi-C contacts at peak pixels was enriched relative to four local neighborhoods (donut, bottom left, horizontal, vertical; see reference 3 for full definitions) and this enrichment was significantly greater than the enrichment at a control set of randomly shuffled loops (as described below) thereby validating our manual loop annotation (Fig. S6). For all subsequent analysis, the central 1 kb of each loop anchor was used.

We also used the HiCCUPS algorithm (3, 31) to annotate chromatin loops. HiCCUPS was run with options -k KR -r 1000 -f 0.001 -p 10 -i 20 -t 0.02,2.5,2.5,2.5 -d 20000. We further filtered the loop list by requiring loops to be greater than 10 kb in size (due to the strong signal along the diagonal, it is hard to unambiguously identify very small loops), to result from collapsing more than 3 enriched pixels to a single peak pixel (to eliminate single, double, or triple pixel blowouts from being called a loop), and to be on chromosome X, 2L, 2R, 3L, 3R, or 4. This resulted in 206 loops. Using the manually annotated loop list as a gold standard (3), HiCCUPS had a 75.6% false positive rate and a 57.5% false negative rate. This elevated error rate is likely due to the fact that HiCCUPS was designed to annotate loops in mammals. Visual inspection of the HiCCUPS-annotated loops identified that the majority of false positives were due to the small feature size of *Drosophila* loops and especially TADs, which made HiCCUPS loop annotation difficult at high resolution and resulted in compartment flips being falsely identified as loops. Due to the extremely high false positive rate we only report results using the manual annotation. However, Pc and Rad21 were also significantly enriched at HiCCUPS-annotated loop anchors (Fig. S7) validating conclusions from our manual annotation. As expected for any annotation with many false positives, the enrichment of Pc and Rad21 at HiCCUPS-annotated loop anchors was less than that for manually-annotated loop anchors.

### Relationships between Loops and TADs

To determine if loops are spatially close to TADs, we determined the Euclidean distance (i.e. the two-dimensional distance between pixels *i*_1_,*j*_1_ and *i*_2_,*j*_2_ in the Hi-C matrix; see reference 8) between a loop (i.e. focal peak of contact enrichment) and the closest TAD corner (smallest Euclidean distance) for all loops. Since TAD sizes differ between flies and humans the Euclidean distance was normalized by the size of the closest TAD for each loop in the respective species.

To determine if loops overlapped TAD corners, that is if loops demarcate TADs, for every loop at location *M_i_,_j_* we determined if there was a TAD corner within distance 0.2*|i-j| of the loop (3). The procedure was repeated for every TAD with TAD corner at *M_i_,_j_* to assess the overlap of TADs with loops.

### Enrichment of Loop Anchors at Non-histone Protein Binding Sites

ChIP-seq reads were mapped to chromosomes X, 2L, 2R, 3L, 3R, and 4 of the dm3/BDGP Release 5 of the *Drosophila melanogaster* genome using bowtie2 with option --very-sensitive. Reads were filtered to include only properly paired reads and reads with a MAPQ >= 30. PCR duplicates were removed with picard. MACS2 (34) was used to call peaks by running macs2 callpeak with parameters --SPMR --keep-dup -g dm -f BAMPE and signal tracks were computed by running macs2 bdgcmp with parameter -m FE. Histone modification ChIP-seq data was processed as above, except the macs2 callpeak parameters --SPMR --keep-dup all --nomodel --nolambda --broad -g dm -f BAMPE were used.

The percentage of unique loop anchors at non-histone protein binding sites was calculated by dividing the number of loop anchors overlapping a ChIP-seq peak by the total number of loop anchors.

For each ChIP-seq dataset we iterated through the list of unique loop anchors counting how many unique loop anchors overlapped with a ChIP-seq peak. We then used bootstrapping to estimate the expected random distribution of counts and to get the significance of enrichment of unique loop anchors at ChIP-seq peaks. This is done by shuffling the order of unique loop anchors while ensuring that anchors remain on their respective chromosome, are not merged in the shuffle, and are excluded from genome assembly gaps. Then, we counted how many shuffled, unique loop anchors overlapped with a ChIP-seq peak. This procedure was repeated 10,000 times generating a new count each time. Enrichments and 95% confidence intervals are then determined by comparing the observed count of unique loop anchors overlapping ChIP-seq peaks to the distribution of shuffled counts.

### Enrichment/Depletion of Loops and Loop Anchors Within Epigenetic Classes

For each epigenetic class as defined in reference 12 we iterated through the list of loops and unique loop anchors counting how many loops/anchors overlapped in their entirety with each epigenetic class. Enrichment/depletion of loops/anchors in each epigenetic class was determined by counting how many unique loops/anchors overlapped in their entirety with an epigenetic class relative to a random shuffle control of loops/anchors using the bootstrapping approach described above, except that a Z-score and p-value for each epigenetic class was used to assess enrichment/depletion by comparing the observed count of unique loops/anchors in each epigenetic class to the distribution of shuffled counts.

The percentage of loops/anchors lying within each epigenetic was calculated by dividing the number of loops/anchors within each epigenetic class by the total number of loops/anchors. Only loops/anchors where the entire loop/anchor overlapped an epigenetic class were considered. Since many loops/anchors overlap two or more epigenetic classes the total percentage of loops/anchors lying within each epigenetic class does not sum to 100%.

### PHO and GAGA Motifs at PRC1 Loop Anchors

PHO and GAGA motif position weight matrices were based on (35). Motifs were identified using STORM (36) and the percentage of PRC1 loop anchors containing both PHO and GAGA motifs was calculated by dividing the number of PRC1 loop anchors overlapping both a PHO and GAGA motif with the total number of PRC1 loop anchors. 10,000 random shuffles of PRC1 loop anchors, similar to that described above, was used as a control. Unique PRC1 loop anchors were identified as those loop anchors overlapping a *Pc* ChIP-seq peak with 30-fold or greater ChIP enrichment.

### Gene Expression at Loop Anchors

RNA-seq data from modENCODE (37) (modENCODE accession modENCODE_4395) was downloaded from release 5.57 of FlyBase (ftp://ftp.flybase.net/releases/FB2014_03/precomputed_files/genes/gene_rpkm_report_fb_2014_03.tsv.gz). RPKM values were extracted for protein-coding genes (13,931 in total) from the Kc167 cell line dataset (FlyBase ID: FBlc0000269).

The percentage of unique PRC1 loop anchors at gene promoters (defined as the set of 1 kb windows centered on the transcription start site for each gene) was calculated by dividing the number of unique PRC1 loop anchors overlapping a gene promoter by the total number of unique PRC1 loop anchors.

Significant differences between RNA-seq RPKM values between PRC1 loop-anchor promoters, promoters bound by PRC1 not at loop anchors, promoters at loop anchors not bound by PRC1, and promoters not at loop anchors and not bound by PRC1 were compared using a one-sided Mann–Whitney *U*-test.

GO term enrichment was performed using DAVID (38) 6.8 Beta (May 2016 knowledgebase) with default parameters. Only GO terms with a Benjamini corrected P-value less than or equal to 10^−3^ were considered. REVIGO (39) was used to remove redundant GO terms with default parameters except a cut-off value (C) of 0.5 was used and the size of the GO term database was set using the *Drosophila melanogaster* GO annotation database. Reduced redundancy GO term analysis is shown in Fig. 3D, complete GO term analysis is shown in Extended Data Table 3. No cellular component GO terms were significantly enriched.

---

**Fig. S1.**
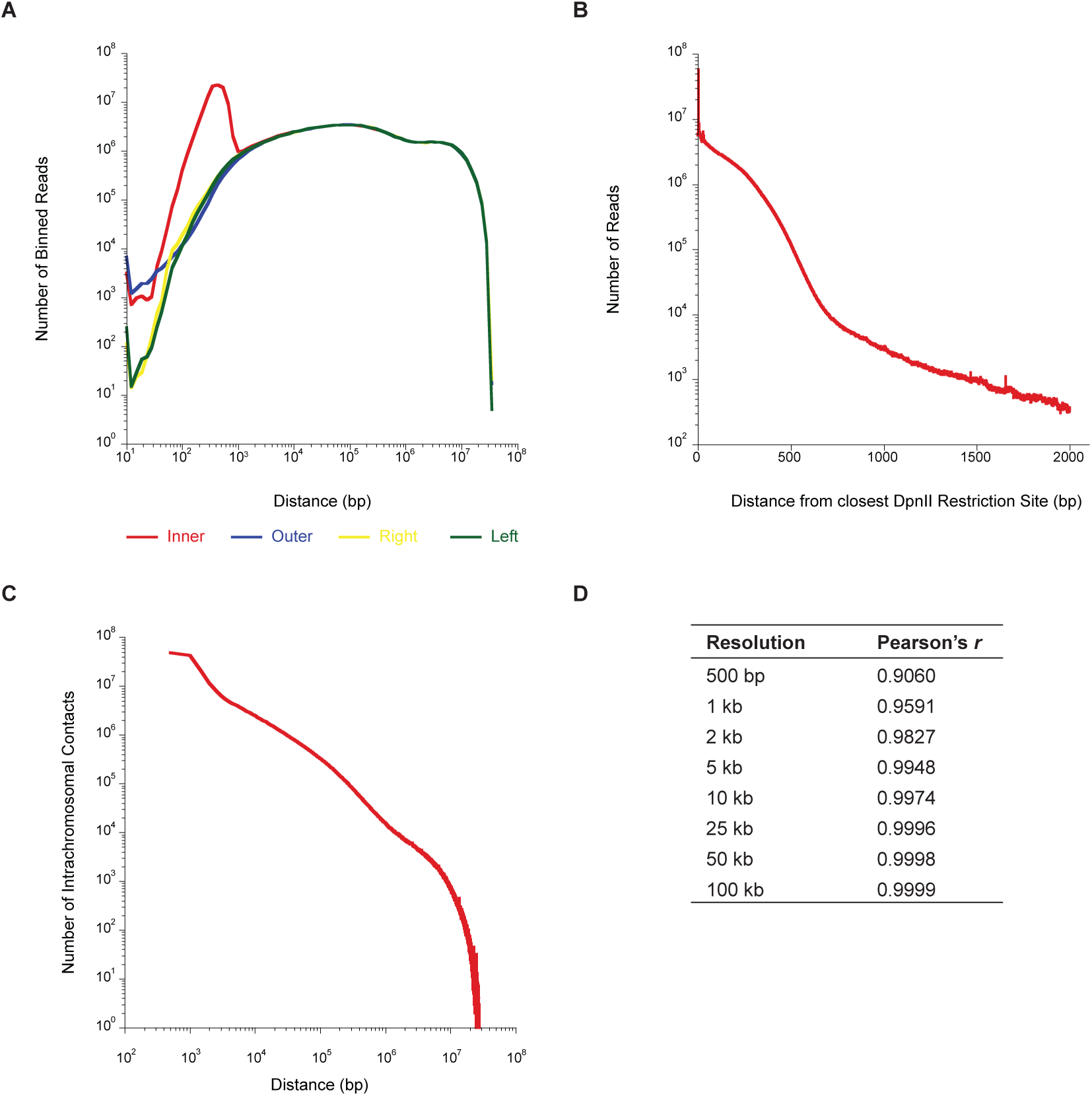
Quality assessment and reproducibility of sub-kilobase resolution *Drosophila* Hi-C. **(A)** Distribution of four read orientations (inner, outer, left, right) as a function of distance between reads. **(B)** Number of reads at the distance indicated on the abscissa to the closest DpnII site. **(C)** Number of intrachromosomal contacts separated by the distance given on the abscissa. **(D)** Pearson’s *r* between biological replicates at the indicated Hi-C contact map resolution. Replicates were highly correlated at all resolutions.

**Fig. S2.**
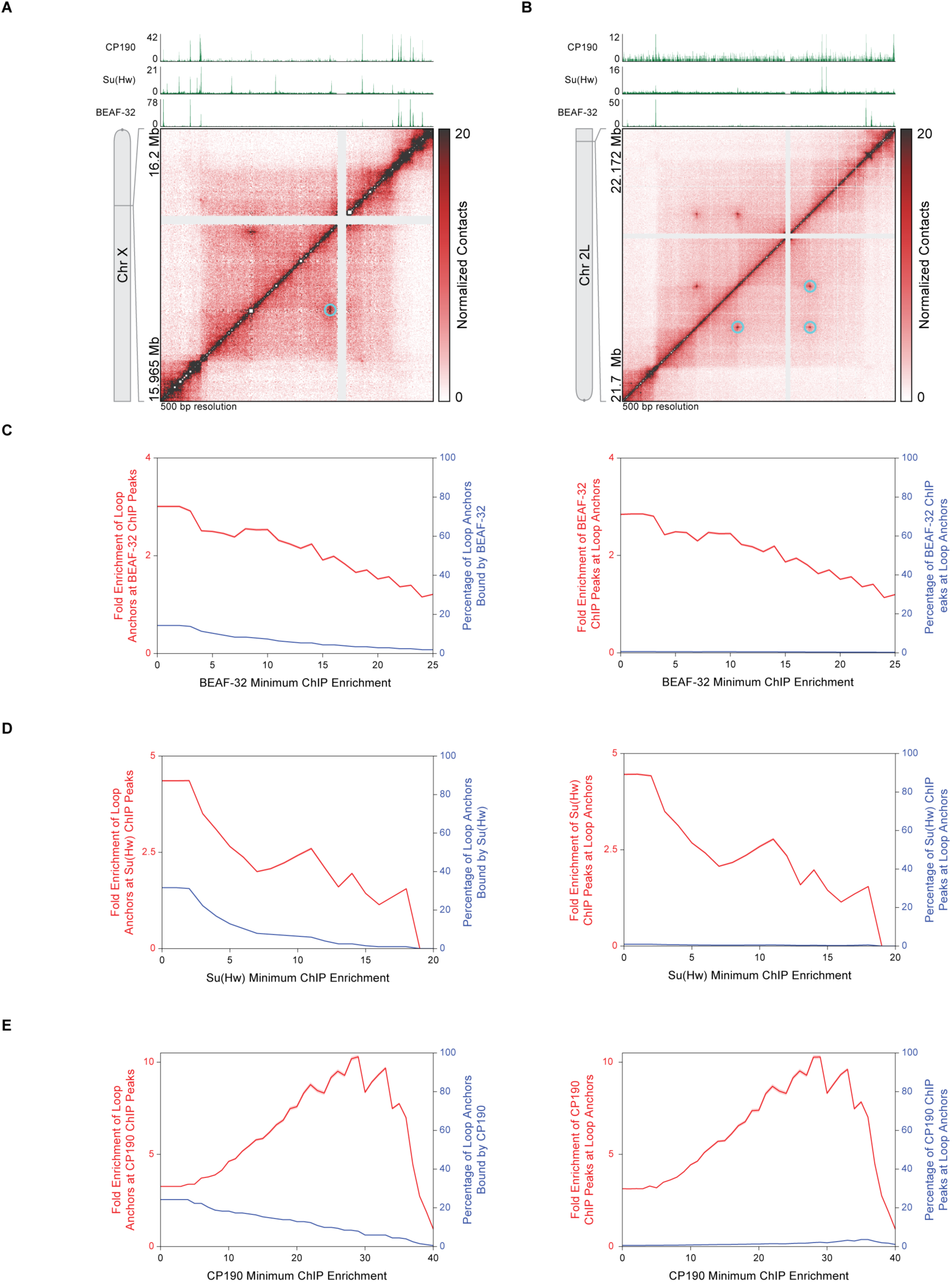
*Drosophila* insulators BEAF-32, Su(Hw), and CP190 are not strongly enriched at loop anchors. **(A)** Hi-C contact map at 500 bp resolution reproduced from Fig. 2A of a region of chromosome X shows chromatin loops (cyan circles). CP190, Su(Hw), and BEAF-32 ChIP-seq profiles are aligned above the map. **(B)** Hi-C contact map at 500 bp resolution reproduced from Fig. 2B of a region of chromosome 2L shows a small network of chromatin loops (cyan circles). CP190, Su(Hw), and BEAF-32 ChIP-seq profiles are aligned above the map. **(C)** Left, fold enrichment of loop anchors at BEAF-32 ChIP peaks (red) and percentage of loop anchors bound by BEAF-32 (blue) at the BEAF-32 minimum ChIP enrichment indicated on the abscissa. Right, fold enrichment of BEAF-32 ChIP peaks at loop anchors (red) and percentage of BEAF-32 ChIP peaks at loop anchors (blue) at the BEAF-32 minimum ChIP enrichment indicated on the abscissa. Red shading indicates 95% confidence interval. **(D)** Same as **(C)** except for Su(Hw). **(E)** Same as **(C)** except for CP190.

**Fig. S3.**
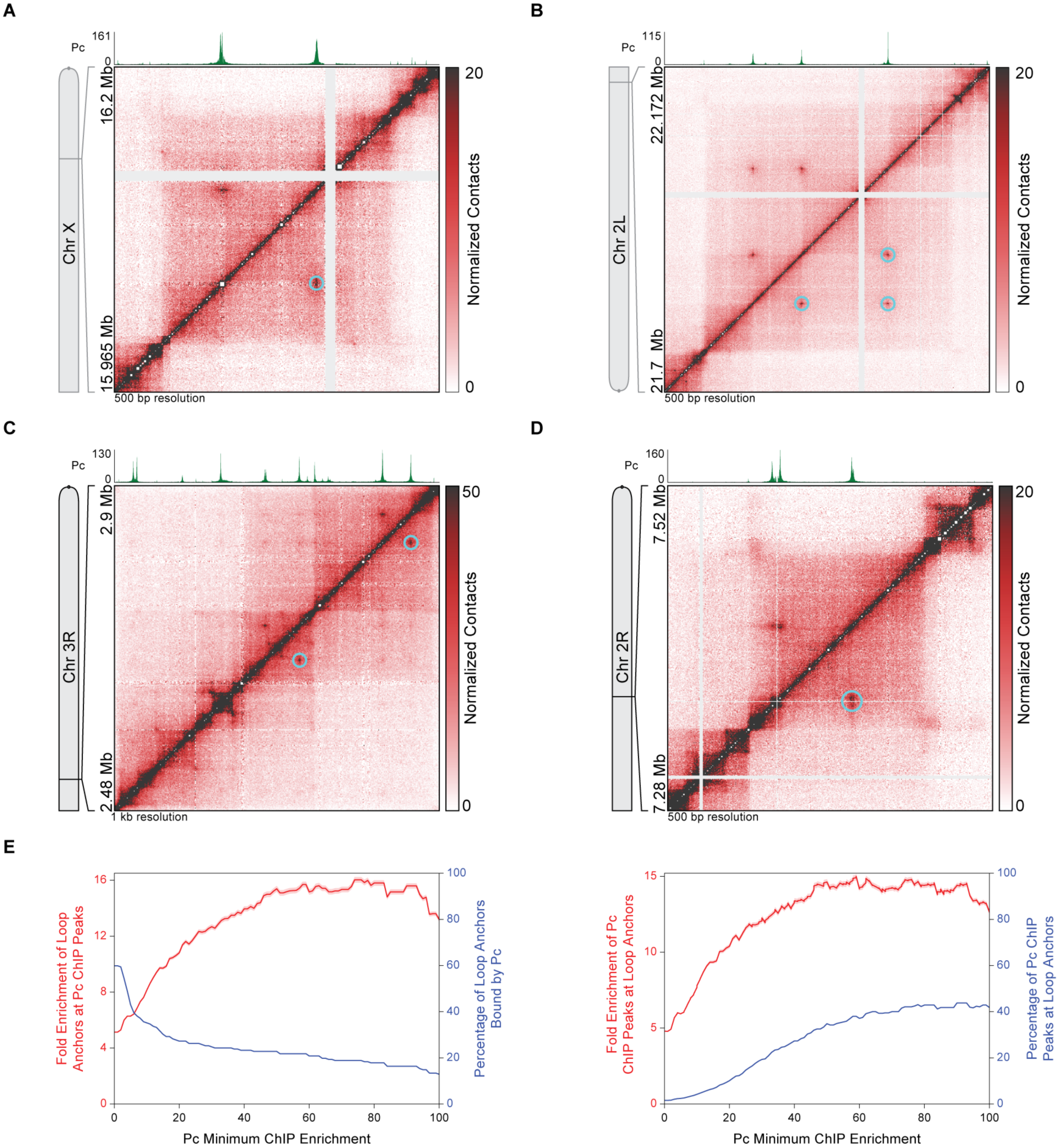
Validation of PRC1 subunit Pc at Loop Anchors. **(A)** Hi-C contact map at 500 bp resolution reproduced from Fig. 2A of a region of chromosome X shows chromatin loops (cyan circles) that align with Pc ChIP-seq peaks. Pc ChIP-seq profile from an antibody different from that in Fig. 2A is aligned above the map. **(B)** Hi-C contact map at 500 bp resolution reproduced from Fig. 2B of a region of chromosome 2L shows a small network of chromatin loops (cyan circles) that align with Pc ChIP-seq peaks. Pc ChIP-seq profile from an antibody different from that in Fig. 2B is aligned above the map. **(C)** Hi-C contact map at 1 kb resolution reproduced from Fig. 3A shows chromatin loops (cyan circles) at the ANT-C Hox gene complex that align with Pc ChIP-seq peaks. Pc ChIP-seq profile from an antibody different from that in Fig. 3A is aligned above the map. **(D)** Hi-C contact map at 500 bp resolution reproduced from Fig. 3B shows chromatin loops (cyan circles) at the *inv* and *en* promoters that align with Pc ChIP-seq peaks. Pc ChIP-seq profile from an antibody different from that in Fig. 3B is aligned above the map. **(E)** Left, fold enrichment of loop anchors at Pc ChIP peaks (red) and percentage of loop anchors bound by Pc (blue) at the Pc minimum ChIP enrichment indicated on the abscissa. Right, fold enrichment of Pc ChIP peaks at loop anchors (red) and percentage of Pc ChIP peaks at loop anchors (blue) at the Pc minimum ChIP enrichment indicated on the abscissa. Pc ChIP peaks are from ChIP-seq data from an antibody different from that in Fig. 2F. Red shading indicates 95% confidence interval.

**Fig. S4.**
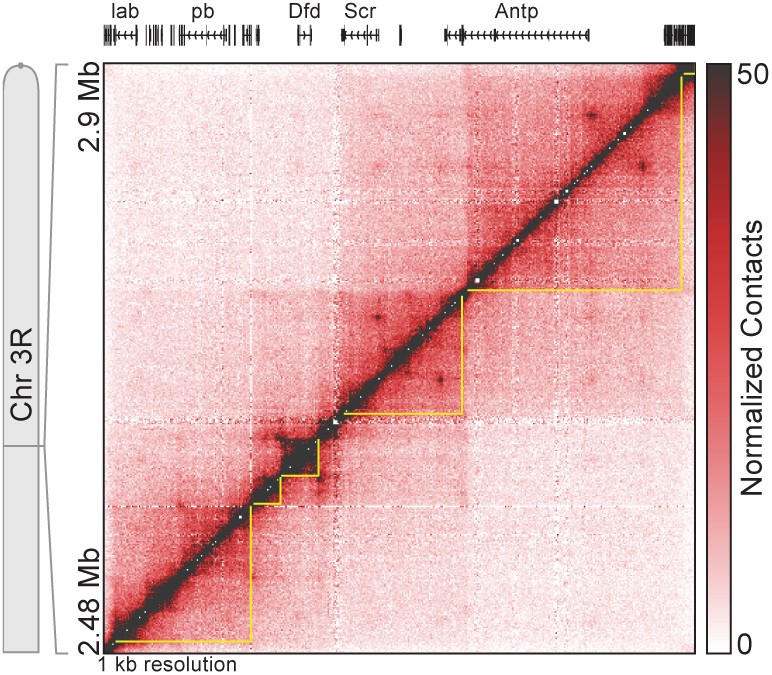
TAD organization at the ANT-C Hox gene complex. Hi-C contact map at 1 kb resolution reproduced from Fig. 3A shows that each TAD (yellow outlines) at the ANT-C Hox gene complex contains one, or, at most, two homeotic gene promoters.

**Fig. S5.**
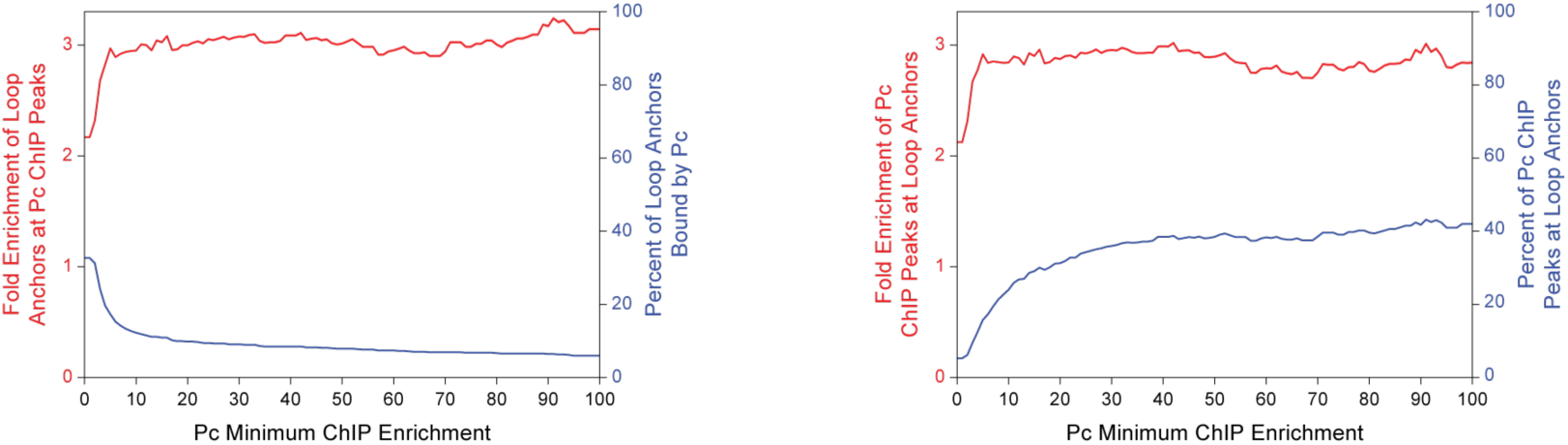
Enrichment of PRC1 subunit Pc at an Independent Set of Loop Anchors. Left, fold enrichment of loop anchors at Pc ChIP peaks (red) and percentage of loop anchors bound by Pc (blue) at the Pc minimum ChIP enrichment indicated on the abscissa. Right, fold enrichment of Pc ChIP peaks at loop anchors (red) and percentage of Pc ChIP peaks at loop anchors (blue) at the Pc minimum ChIP enrichment indicated on the abscissa. Loop anchors are from reference 26 and were converted from the dm6 to dm3 genome assembly using the FlyBase *Drosophila* Sequence Coordinates Converter. Red shading indicates 95% confidence interval.

**Fig. S6.**
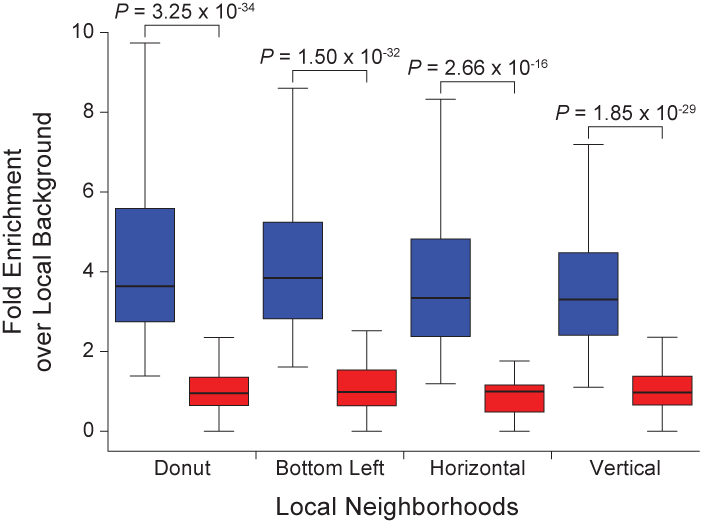
Enrichment of Hi-C Contacts at Loops. Box plots of the number of Hi-C contacts at manually annotated loops compared to the number of Hi-C contacts in four local neighborhoods (donut, bottom left, horizontal, vertical; see reference (3) for full definitions). Enrichment at manually annotated loops was significantly greater than the enrichment at a control set of randomly shuffled loops. Significance computed using a one-sided Mann–Whitney *U*-test.

**Fig. S7.**
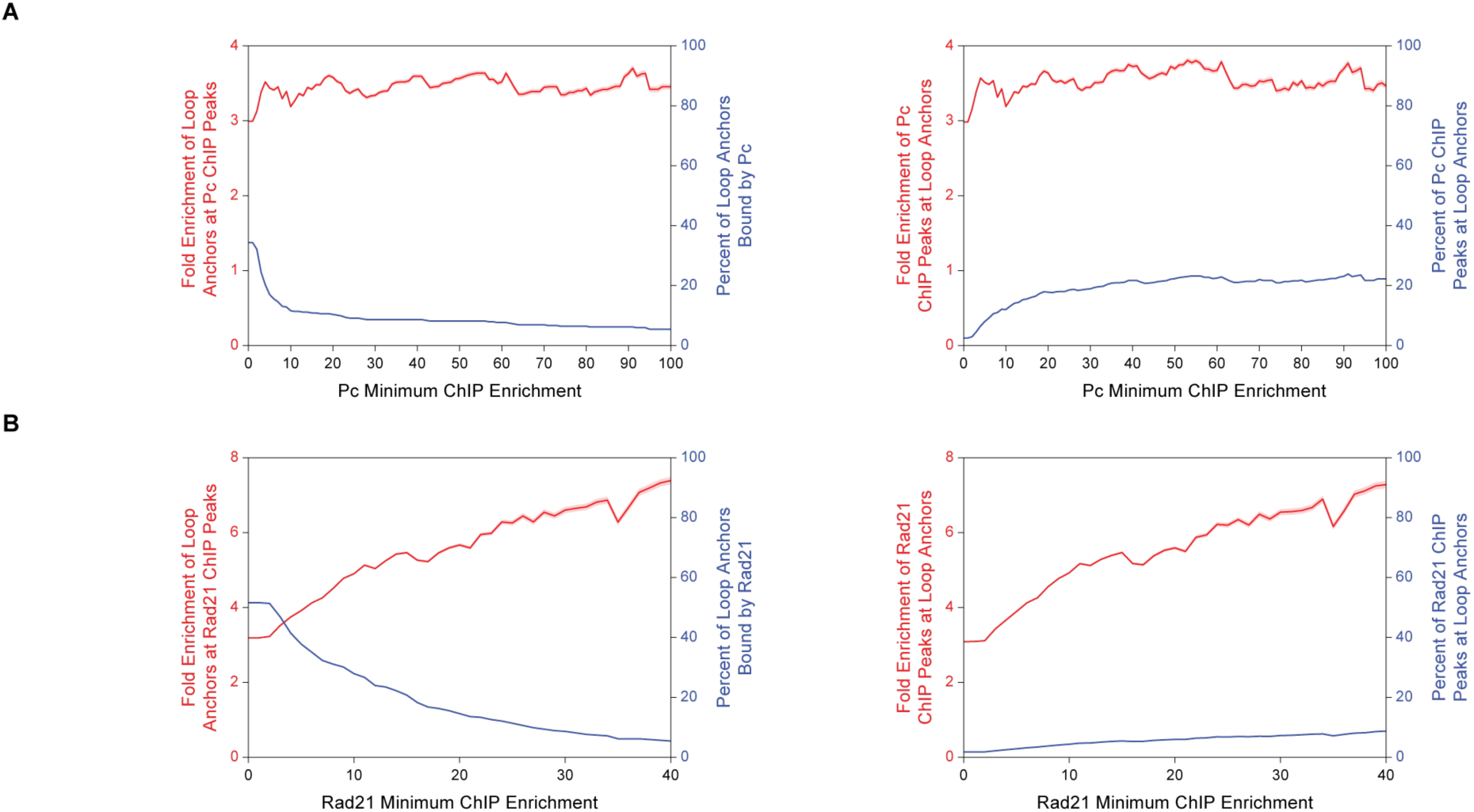
Enrichment of Pc and Rad21 at HiCCUPS-Annotated Loop Anchors. **(A)** Left, fold enrichment of loop anchors at Pc ChIP peaks (red) and percentage of loop anchors bound by Pc (blue) at the Pc minimum ChIP enrichment indicated on the abscissa. Right, fold enrichment of Pc ChIP peaks at loop anchors (red) and percentage of Pc ChIP peaks at loop anchors (blue) at the Pc minimum ChIP enrichment indicated on the abscissa. Loop anchors are from loops identified by HiCCUPS (see *SI Appendix, Supplementary Materials and Methods).* Red shading indicates 95% confidence interval. **(B)** Same as **(A)** except for Rad21.

**Table S1.** Hi-C sequencing statistics and quality metrics.

|  | Primary | Replicate | Merged |
| --- | --- | --- | --- |
| **Sequencing Statistics** |  |  |  |
| Sequenced Reads | 532,673,670 | 554,361,974 | 1,087,035,644 |
| **Alignment Statistics\*** |  |  |  |
| Normal Paired | 395,699,521 (74.29%) | 408,451,484 (73.68%) | 804,151,005 (73.98%) |
| Chimeric Paired | 87,524,998 (16.43%) | 92,918,735 (16.76%) | 180,443,733 (16.60%) |
| Unmapped | 10,388,903 (1.95%) | 10,203,756 (1.84%) | 20,592,659 (1.89%) |
| Ligation Motif Present | 247,207,877 (46.41%) | 264,925,937 (47.79%) | 512,133,814 (47.11%) |
| **Complexity Statistics\*** |  |  |  |
| Alignable (Normal+Chimeric) | 483,224,519 (90.72%) | 501,370,219 (90.44%) | 984,594,738 (90.58%) |
| Unique Reads | 408,175,481 (76.63%) | 424,129,873 (76.51%) | 832,305,354 (76.57%) |
| PCR Duplicates | 74,802,581 (14.04%) | 77,032,979 (13.90%) | 151,835,560 (13.97%) |
| Optical Duplicates | 246,457 (0.05%) | 207,367 (0.04%) | 453,824 (0.04%) |
| **Unique Read Statistics\*\*** |  |  |  |
| Intra-fragment Reads | 25,145,283<br>(4.72% / 6.16%) | 25,523,374<br>(4.60% / 6.02%) | 50,668,657<br>(4.66% / 6.09%) |
| Below MAPQ30 | 123,173,623<br>(23.12% / 30.18%) | 129,753,530<br>(23.41% / 30.59%) | 252,927,153<br>(23.27% / 30.39%) |
| **Hi-C Read Statistics\*\*** |  |  |  |
| Hi-C Contacts | 259,856,575<br>(48.78% / 63.66%) | 268,852,969<br>(48.50% / 63.39%) | 528,709,544<br>(48.64% / 63.52%) |
| Ligation Motif Present | 95,704,474<br>(17.97% / 23.45%) | 101,798,673<br>(18.36% / 24.00%) | 197,503,147<br>(18.17% / 23.73%) |
| 3' Bias (Long Range) | 69% - 31% | 70% - 30% | 70% - 30% |
| Pair Type % (L-I-O-R) | 25% - 25% - 25% - 25% | 25% - 25% - 25% - 25% | 25% - 25% - 25% - 25% |
| Inter-chromosomal | 12,056,707<br>(2.26% / 2.95%) | 12,038,653<br>(2.17% / 2.84%) | 24,095,360<br>(2.22% / 2.90%) |
| Intra-chromosomal | 247,799,868<br>(46.52% / 60.71%) | 256,814,316<br>(46.33% / 60.55%) | 504,614,184<br>(46.42% / 60.63%) |
| Short Range (< 20 kb) | 107,655,986<br>(20.21% / 26.37%) | 114,484,535<br>(20.65% / 26.99%) | 222,140,521<br>(20.44% / 26.69%) |
| Long Range (> 20 kb) | 140,138,997<br>(26.31% / 34.33%) | 142,325,315<br>(25.67% / 33.56%) | 282,464,312<br>(25.98% / 33.94%) |

\* % Sequenced Reads
† % Sequenced Reads / % Unique Reads
For definitions of each metric see reference 3.

**Table S2. Kc167 TAD coordinates.**

Juicebox format. dm3 genome assembly.

See attached Excel file.

**Table S3. Kc167 loop coordinates.**

Table S3. Kc167 loop coordinates.

Juicebox format. dm3 genome assembly.

See attached Excel file.

**Table S4. Complete GO terms for PRC1 loop anchors.**

See attached Excel file.

**Table S5.** External datasets used in this study.

| **External Dataset** | **Reference** | **Accession Number** | **Antibody (if given/applicable)** | **Notes** |
| --- | --- | --- | --- | --- |
| Kc167 5 chromatin classes | 12 | GSE22069 |  |  |
| Kc167 BEAF-32 ChIP-Seq | 11 | GSE30740 |  |  |
| Kc167 CP190 ChIP-Seq | 11 | GSE30740 |  |  |
| Kc167 CTCF ChIP-Seq | 11 | GSE30740 |  |  |
| Kc167 H3K27me3 ChIP-Seq | 14 | GSE37444 | Millipore Cat. # 07-449 Lot # JBC1924326 |  |
| Kc167 Pc ChIP-Seq | 15 | GSE63518 | RJ | Fig. 2, 3, S5, S7 |
| Kc167 Pc ChIP-Seq | 15 | GSE63518 | VP | Fig. S3 |
| Kc167 RNA-seq | 37 | modENCODE_4395 |  |  |
| Kc167 Su(Hw) ChIP-Seq | 11 | GSE30740 |  |  |
| Kc167 loops (independently annotated) | 26 | GSE80702 |  | Fig. S5 |
| GM12878 Hi-C loops | 3 | GSE63525 |  |  |
| GM12878 Hi-C TADs (contact domains) | 3 | GSE63525 |  |  |

